# Development of an Ectopic huLiver Model for *Plasmodium* Liver Stage Infection

**DOI:** 10.1101/2022.12.01.518796

**Authors:** Gabriela Samayoa Reyes, Siobhan Flaherty, Kristina S. Wickham, Sara Viera-Morilla, Pamela Strauch, Alison Roth, Laura Padrón, Conner Jackson, Patricia Meireles, David Calvo, Wanlapa Roobsoong, Niwat Kangwanrangsan, Jetsumon Sattabongkot, Gregory Reichard, Maria José Lafuente-Monasterio, Rosemary Rochford

## Abstract

Early *Plasmodium falciparum* and *P. vivax* infection requires parasite replication within host hepatocytes, referred to as liver stage (LS). However, limited understanding of infection dynamics in human LS exists due to species-specificity challenges. Reported here is a reproducible, easy-to-manipulate, and moderate-cost *in vivo* model to study human Plasmodium LS in mice; the ectopic huLiver model. Ectopic huLiver tumors were generated through subcutaneous injection of the HC-04 cell line and shown to be infectible by both freshly dissected sporozoites and through the bite of infected mosquitoes. Evidence for complete LS development was supported by the transition to blood-stage infection in mice engrafted with human erythrocytes. Additionally, this model was successfully evaluated for its utility in testing antimalarial therapeutics, as supported by primaquine acting as a causal prophylactic against *P. falciparum.* Presented here is a new platform for the study of human *Plasmodium* infection with the potential to aid in drug discovery.

## Introduction

Malaria has a substantial impact on global public health, with an estimated 229 million cases occurring in 2019, and is considered not only the most lethal parasitic infection, but also one of the most prevalent infectious diseases worldwide [1]. The intracellular protozoan parasite of the genus *Plasmodium* is the causative agent, with the species *Plasmodium falciparum* contributing to the majority of malaria-associated morbidity and mortality in humans [2]. The second most prevalent species, *P. vivax,* while less associated with mortality, is the most widely distributed with an estimated one-third of the world’s population at risk of infection [3]. Parasite transmission occurs through the bite of an infected female *Anopheles* mosquito, subsequently sporozoites injected by the mosquito into the dermis travel through the circulatory system until reaching the liver. Within the liver, sporozoites traverse numerous hepatocytes before selecting a single one to invade and form a parasitophorous vacuole (PV) for replication [4]. This initiates the first stage of *Plasmodium* infection called the liver stage (LS). Completion of LS infection results in the formation of thousands of merozoites from a single hepatocyte, which egress and proceed to infect erythrocytes. The invasion of erythrocytes constitutes the second stage of infection, the blood-stage, where the clinical presentation of the disease develops [2,5,6]. While the described processes are representative of *P. falciparum* LS infection, *P. vivax* infection follows a similar pattern. However, *P. vivax* uniquely features dormant LS parasites, hypnozoites, which are the source of the characteristic long-latency and relapse seen after primary infection [7]. Studying LS infection of both *P. falciparum* and *P. vivax* has proved challenging with the limited models available.

Most of the current knowledge about *Plasmodium* LS infection has been generated using murine malaria models; they have provided information from development and host-parasite interactions, to merozoite release from infected hepatocytes. However, there are clear differences between the LS biology of rodent *Plasmodium* species, *P. yoelii* and *P. berghei*, and the human-specific strains, *P. falciparum* and *P. vivax* [2]. One of the main differences is the kinetics of LS infection. Rodent *Plasmodium* LS infection lasts approximately 2 days from infection to release of merozoites, while in humans the *P. falciparum* and *P. vivax* LS infection can take approximately 1 week, and in the case of *P. vivax*, relapse may occur months or even years after the primary infection [8]. Another factor to consider are genes which could have different biological functions in mouse models as compared to humans [9, 10], and that not all proteins expressed by human *Plasmodium* species have orthologs in rodent malaria models [8, 10]. This emphasizes the need for a robust humanized *in vivo* LS infection model, which would ideally support both *P. falciparum* and *P. vivax*.

In 2006, the HC-04 cell line was reported to be the first human hepatocyte (huHep) cell line that supports complete *P. falciparum* and *P. vivax* exo-erythrocytic development, both *Plasmodium* species showed full maturation and release of infective merozoites [11]. The HC-04 cell line, when infected, displays a time-dependent increase in *P. falciparum* 18S gene copy number and *P. vivax* 18S rRNA copy number [12], an indication that both can develop successfully. The infection rate is approximately 0.066% for *P. falciparum,* and 0.041% for *P. vivax* [11].

Previously, complete development of *P. falciparum* LS and liver-to-blood stage transmission has been demonstrated *in vivo* using SCID *Alb-uPA* immunocompromised mice engrafted with huHeps [13], although limitations have been reported [8]. To overcome the deficiencies associated with the SCID *Alb-uPA* model, the immunocompromised fumarylacetoacetate hydrolase–deficient mouse (*Fah^−/−^*, *Rag2^−/−^*, *Il2rg^−/−^*), known as the FRG KO mouse model, was developed. This model when backcrossed to NOD (FRGN KO huHep) can be repopulated with huRBCs, which can support blood-stage infection of both *P. falciparum* and *P. vivax* [8,14–16]. The FRG KO huHep model allows for complete *P. falciparum* LS development and full maturation, with merozoite formation and release and also supports *P. vivax* liver stage development, including hypnozoite development [15]. Although the FRG KO huHep mice can be infected with both *P. falciparum* and *P. vivax* and have proven to be a great tool for LS studies, the model presents several limitations, including the variable cytochrome p450 (CYP) 2D6 metabolic activity, which correlates to the level of repopulation of the mouse liver with human hepatocytes and is subject to donor variability [17, 18]. This affects the overall efficacy of the model for antimalarial therapeutic development where CYP 2D6 activity is required for compound metabolism [17,19–21]. Despite the noted limitations, low-throughput, and high cost, the FRG-huHep model is the best option thus far for studying *Plasmodium* LS development, highlighting the need for additional humanized mouse models.

To address the challenges of previous *Plasmodium* LS *in vivo* models, we developed the ectopic human liver mouse model (ectopic huLiver). The ectopic huLiver model is a tumor model where tumors are formed in extremely immunodeficient NOD/SCID/ *IL2rg^null^* (NSG) mice following engraftment with the HC-04 cell line, and can be humanized by engraftment with huRBCs allowing for transition to blood-stage infection. Here we present data on the evaluation of infection of the ectopic huLiver with *P. falciparum* and *P. vivax* and the potential for this model as a tool in drug discovery.

## Materials and Methods

### Experimental model and details

All animal experiments carried out at the University of Colorado were approved by Institutional Animal Care and Use Committee (IACUC) at University of Colorado, Anschutz Medical Campus, and all experimental methods were carried out in accordance with the approved guidelines.

All studies were conducted in accordance with the GSK Policy on the Care, Welfare and Treatment of Laboratory Animals and were reviewed the Institutional Animal Care and Use Committee either at GSK or by the ethical review process at the institution where the work was performed

#### Mice

Female 6-week old immunocompromised NOD/SCID/ *IL2rg^null^* (NSG) mice were purchased from Jackson Laboratories (Bar Harbor, ME USA). Mice were housed in groups of up to 5 mice per cage. Experimental groups were based on size of the tumor, to include mice with similar sized tumors in both groups.

Female 6-week FRG-huHep (on C57/ black 6 background) [22] and FRG NOD huHep mice with human chimeric livers were purchased from the Yecuris Corporation (Tigard, OR).

#### huRBCs

For experiments that required huRBC engraftment, blood from incomplete donations, or erythrocyte concentrates of malaria-negative donors were generously provided by the Spanish Red Cross blood bank in Madrid, Spain. The human biological samples were sourced ethically and their research use was in accordance with the terms of informed consents under an IRB/EC approved protocol.

#### HC-04 cell line

HC-04 human hepatocytes derived from the liver of a hepatoma patient were obtained from BEI Resources, NAID, NIH (deposited by J. Sattabongkot, catalog no. MRA0975). Cells were cultured in in Nutrient Mixture (HyClone, Marlborough, MA) with Modified Eagle Media (Gibco, Waltham, MA), containing 10% FBS, Pen-Strep (Gibco), HEPES (Gibco), and Sodium Bicarbonate (Corning, Corning, NY), L-Glutamine (Gibco) and maintained at 37°C with 5% CO2. Cells were split every 3-days or when confluency of 80% was reached.

### Method details

#### Mice

##### NSG mice

7-8-week-old mice were subcutaneously (s.c.) injected in the flank with 5.0 × 10^6^ HC-04 cells or HC-04 EphA2^High^ CD81 ^High^, resuspended in MEM medium. Ectopic huLiver growth was monitored daily using caliper measures until reaching a volume of .5 cm^3^ approximately 14-19 days post tumor engraftment, at this point the tissue was considered infectible with human *Plasmodium* species. Ectopic huLiver volume was calculated using the formula ½ (length * width^2^).

##### FRG-huHep mice

human serum albumin levels, blood samples from each repopulated mouse were collected via the saphenous vein and analyzed using the Human Albumin ELISA Quantitation Set (Abcam, Cambridge, UK) according to the manufacture’s protocol. All mice within the same experiment were engrafted with primary human hepatocytes originating from the same donor selected from an assortment of prequalified lots. Mice were cycled on 2-(2-nitro-4-trifluoromethylbenzoyl)-1, 3-cyclohexanedione (NTBC) on a 4-week schedule. In week 1, mouse water was supplemented with NTBC at 16 mg/L for 3 days [8]. The drug was then removed for week 2 [8]. This schedule continued until the end of the experiment. Animals did not receive NTBC 3 days prior to experiments.

##### Experimental endpoints

were based on tumor size greater than 10% body weight or exceeding 20 mm in any dimension, tumors that ulcerate and/or become necrotic. If mouse was otherwise healthy it was kept until data collection day according experimental plan. Animals were euthanized by carbon dioxide asphyxiation followed by cervical dislocation.

#### Ectopic huLiver characterization

##### Immunofluorescence staining of DNA and Laminin

Uninfected ectopic huLiver tissue was embedded in Tissue-Tek OCT (Sakura Finetek, Torrance, CA), snap frozen on dry ice and stored at −80°C. Tissue was sectioned using a cryostat (CryoJane adaptation, Leica) into 6 μm thick sections and transferred onto glass slides. Slides were dried at RT for 30 min, followed by fixation for 30 sec in 50% acetone and then 3 min in 100% acetone. Tissue sections were rehydrated in PBS for 20 min, blocked with goat serum and anti-mouse unlabeled IgG, in a 2% BSA/0.05% Tween20/PBS solution for 15 min at RT. Slides were washed 3x with PBS, then stained with a polyclonal rabbit anti-laminin Dylight 488 antibody (Invitrogen, Carlsbad, CA) (1:100) for 45 min at RT. Slides were washed 3x with PBS, followed by the addition of DAPI ProLong mounting medium (Invitrogen, Carlsbad, CA) and stored at RT for up to 24 hours image capture.

##### *Plasmodium* sporozoite production in *Anopheles* mosquitoes

In vitro P. falciparum Nijmegen falciparum strain 54 (NF54HT) [23] and NF54HT-GFP-luc parasite culture, which constitutively expresses luciferase under the EF1α promoter [24] were obtained from the Malaria Research and Reference Reagent Resource [25]. All *Plasmodium* strains were grown according to the method previously described by Trager and Jensen [26]. Mature sexual-stage gametocytes were induced as previously described [8] and fed to female *Anopheles stephensi* mosquitoes to initiate infection. *An. stephensi* colony was established from eggs kindly supplied by Michael Delves and Mark Tunncliffe from Imperial College, London in 2014. Generated as previously described [27]. From March 2014 until date, *An. stephensi* line SDa 500 mosquitoes have been rearing at at R&D of GlaxoSmithKline in Madrid, Spain. GSK breeds [28]. Larval stages were reared following standard protocols as described in the MR4 manual [8]. Adult female mosquitoes were infected 3 to 5 days after emergence by feeding with *Plasmodium* infected mice. Infection progression was monitored at 7 days post-feeding by counting oocysts on the midgut of infected mosquitoes using the mercurochrome staining method [28, 29].

#### P. falciparum infections

For mosquito bite infection, FRG-huHep mice and ectopic huLiver were anesthetized with isoflurane, the mice were placed in the induction chamber and the gas turned on at 2-3% with a flow rate of 0.8-1.0 liter/min for 7-10 minutes. Once anesthetize the mice were placed on the top of a mosquito cup containing 20 infected mosquitoes for 20 min. Mice were then transferred to a second cup, for a second infection round [30]. FRG-huHep mice were positioned on their ventral side, and the ectopic huLiver mice were positioned laterally and shaved to allow mosquito access to the ectopic huLiver during the mosquito bite challenge.

For administration of sporozoites by intravenous (i.v) injection into the tail vein (FRG-huHep mice) or intratumorally (i.t) (ectopic huLiver mice), sporozoites were obtained from 14- to 19-day-old mosquito salivary glands that were dissected, and resuspended in RPMI medium according to previously published methods [30]. In brief, salivary glands of 20 infected mosquitoes were dissected, placed in 50 μl of ice-cold RPMI 1640 medium (Gibco, Grand Island, NY, USA), pH 8.2 and ground with a sterile pestle [31]. The homogenate obtained was cleared by centrifugation for 5 min at 1,200 rpm. The released sporozoites were counted in a hemocytometer and kept on ice until used, but for no more than 1 hour to avoid a reduction in parasite infectivity [28, 32].

#### Engraftment with huRBCs

Prior to injection, blood was stored at 4°C and washed 2x with RPMI 1640, 25 mM HEPES (Sigma-Aldrich, St. Louis, MO) containing 7×10^−3^ mM hypoxanthine (Sigma-Aldrich, St. Louis, MO) at RT. The buffy coat was removed by aspiration with erythrocytes resuspended at 50% hematocrit in RPMI 1640, 25% decomplemented human serum (Sigma-Aldrich, St. Louis, MO), and 3.1 mM hypoxanthine. huRBC suspension was warmed at 37°C for 20 min prior to intraperitoneal (i.p.) injection. Mice were i.p injected with 1 ml of huRBCs containing 70% O^+^ human erythrocytes in RPMI 1640 (25 mM HEPES, 2 mM L-glutamine) supplemented with 50 μM hypoxanthine plus 10% human serum. Engraftment continued daily using 1 ml huRBCs throughout the experiment, in order to reach the minimum desired percentage of huRBCs ≥ 70% (measured by flow cytometry as previously described [33]) by 7 days post mosquito bite challenge.

#### Quantification of liver stage parasite burden

##### LS-to-blood-stage transition determination using P. falciparum 18S qRT-PCR

Starting at 9 days post-infection blood samples, of both FRG-huHep and ectopic huLiver, were collected by submandibular bleeding [34] and mixed 1:10 (v/v) with heparin. The sample was either processed within 2 hours of collection, to avoid RNA degradation or frozen in lysis buffer supplemented with β-mercaptoethanol until processing. Whole blood RNA was extracted using the GeneJET Whole Blood RNA Purification Kit (Thermo Fisher, Waltham, MA), following the manufacturer’s protocol. The isolated RNA was quantified using a NanoDrop spectrophotometer (Thermo Fisher, Waltham, MA). RNA was reverse transcribed using the High-Capacity cDNA Reverse Transcription Kit (Applied Biosystems, Foster City, CA). The resulting cDNA was used for the amplification *of P. falciparum* 18S rRNA and the housekeeping gene Glyceraldehyde-3-Phosphate Dehydrogenase (hu*GAPDH)*. The primer set and conditions used to detect *P. falciparum* 18S have been previously described [35]. Briefly, 2 μL of cDNA were used in a reaction mixture with 150 nM of each primer and 250 nM of the probe and 1X TaqMan Fast Advanced Master Mix (Applied Biosystems, Foster City, CA). Reactions were run in a ViiA 7 Real-Time PCR system (Applied Biosystems, Foster City, CA) using fast reaction presets. Reverse transcriptase negative and non-template controls were used. Parasite 18S copies were quantified by generating a standard curve with a plasmid control. The quality of all samples was verified by amplification of human *GAPDH*.

##### Immunofluorescence staining of MSP1 and EBA-175

Immunofluorescent (IF) staining was performed on infected ectopic huLiver tissue for detection of *P.falciparum* MSP1 (kindly provided by Dr. A. Evelina, Walter Reed Army Institute (WRAIR)) and EBA-175. The reagent EBA-175 was obtained through BEI Resources, NIAID, NIH: Monoclonal Antibody R217 Anti-*Plasmodium falciparum* Erythrocyte Binding Antigen-175 RII (produced *in vitro*), MRA-711A, contributed by B. Kim Lee Sim and NIAID/NIH. Ectopic huLiver tissue was processed and stained using the methods described above with polyclonal rabbit anti-MSP1 (1:100) and monoclonal mouse anti-EBA-175 (1:100) followed with goat anti-rabbit IgG (1:100) (Abcam, Cambridge, UK), and goat α-mouse IgG1 (1:100) (Abcam, Cambridge, UK).

#### Characterizing ectopic huLiver metabolism and evaluating primaquine as a prophylactic in the ectopic huLiver model

##### Immunohistochemistry staining for ASGPR1

Uninfected ectopic liver tissue was fixed in a 10% buffered formalin solution, transferred to a 70% ethanol solution after 24 hours, paraffin embedded and sectioned into 6 μm thick sections using a cryostat. To determine expression levels of ASGPR1, anti-ASGPR1 IHC staining was performed as previously described [36]. Briefly, slides were deparaffinized and rehydrated with Xylene 2X, followed by a 100% to 70% ethanol gradient, and PBS. Rehydrated sections were antigen retrieved with citrate buffer at 103°C for 10 min. The sections were then incubated with 3% H_2_O_2_ in methanol for 10 min at RT, washed 3x with PBS, and then incubated for 30 min with 10% BSA in a humidified chamber at RT. The primary antibody, rabbit anti-ASGPR1 (Sigma-Aldrich, St. Louis, MO) at 1:500 was applied to the sections and incubated for 2 hours at RT. Isotype control was stained with rabbit IgG (1:500). Sections were then rinsed in PBS and incubated with an HRP-conjugated anti-rabbit secondary antibody from the ImmPress anti-rabbit kit, (Vector Laboratories, Burlingame, CA) for 30 min. Samples were washed and incubated with ImmPact DAB (Vector Laboratories, Burlingame, CA). Sections were further counterstained with Hematoxylin QS then dehydrated and mounted with Vectamount Mounting Media (Vector Laboratories, Burlingame, CA).

##### HC-04 CYP 2D6 expression quantification

RNA was extracted from the HC-04 cell line using the RNeasy Plus Mini Kit (Qiagen, Hilden, Germany), following the manufacturer’s protocol. The isolated RNA was quantified using a NanoDrop spectrophotometer (Thermo Fisher, Waltham MA). HC-04 cell line CYP 2D6 RNA levels were compared against commercially available human liver tissue CYP 2D6 RNA levels (OriGene Technologies, Rockville, MD). RNA was reverse transcribed using the QuantiTect Reverse Transcription Kit (Qiagen, Hilden, Germany), with the resulting cDNA used for amplification of CYP 2D6 rRNA and the housekeeping gene beta-2-microglobulin (B2M). RT-PCR was performed using an iCycler thermocycler equipped with an iCycler iQ real-time PCR detection system (Bio-Rad Laboratories, Hercules, CA). CYP 2D6 primers sequences were used based on previously published data [37] and adjusted as necessary based on NCBI BLAST analysis. RT-PCR was preformed using SYBR Green Master Mix (Bio-Rad Laboratories, Hercules, CA) with the following primers: B2M F: 5’-GAG TAT GCC TGC CGT GTG-3’, B2M R: 5’-AAT CCA AAT GCG GCA TCT-3’, CYP2D6 F: 5’-CGC ATC CCT AAG GGA ACG A-3’ and CYP 2D6 R: 5’-TGC CAG ACG GCC TCA TCC T-3’. The PCR protocol was as follows: 10 min at 95°C, 40 cycles of 15 sec at 95°C and 1 min at 60°C, followed by a denaturation melt curve step. Expression levels were normalized by using the B2M housekeeping gene. Delta-delta cycle threshold (ΔΔC_T_) analysis was used to evaluate CYP 2D6 transcript expression relative to B2M expression in the HC-04 cell line compared with human liver tissue RNA. The formula 2^−ΔΔCt^ was then applied to give the expression fold change between the two samples.

##### Pharmacokinetics of the anti-malarial compound primaquine in ectopic huLiver mice

8 NSG mice with ectopic huLivers were administered 30 mg/kg of primaquine (PQ) with 3 additional mice reserved as controls and dosed with PBS only. For PQ, mice were sacrificed at 30 min and 120 min post first dose and blood was collected by cardiac puncture into a CPDA coated syringe. Ectopic huLiver and mouse liver tissues were harvested and blood was collected, ectopic livers, and mouse livers were snap frozen until PK analysis could be conducted. Samples were extracted and analyzed as described below.

##### Sample preparation and extraction

Standard curves and quality control (QC) samples were prepared by spiking mouse whole blood with PQ, carboxy-PQ, and WR911000 and then performing serial dilutions to desired concentrations. Aliquots of 15 µL of the spiked mouse whole blood from the standard curve and QC samples were spotted on Whatman FTA DMPK-C cards and were allowed to dry at 27°C. Two 3-mm cores were punched from the center of the 15 µL dried blood spot for extraction and placed into a clean Eppendorf tube. The 3-mm cores were submerged in 50 µL of methanol containing internal standard and vortexed gently for 15 min. The samples were then centrifuged at 16,000x*g* for 15 min. The resulting supernatant was transferred to total recovery vials (Waters Corporation, Milford, MA) for liquid chromatography-tandem mass spectrometry (LC-MS/MS) analysis.

##### Instrumentation for PQ and WR compound analysis

An AB Sciex 4000 QTrap triple quadrupole instrument (AB Sciex, Framingham, MA) with an electrospray interface in positive ionization mode was used for the multiple-reaction monitoring (MRM) LC-MS/MS analysis of all the samples. Electrospray ionization conditions were optimized for PQ, carboxy PQ, and WR911000. Separations were achieved on a reversed-phase liquid chromatography Waters XterraC18 column (2.1 by 50 mm; 3.5µm) under ambient (25±2°C) conditions. The mobile phase consisted of solvent A, water with 0.1% formic acid, and solvent B, methanol with 0.1% formic acid. A 6.0-min gradient elution of the analytes was performed at a flow rate of 400 µL/min. PQ, carboxy-PQ, and WR911000, concentrations were interpolated from a combined standard curve.

##### P. falciparum infected ectopic huLiver sensitivity to primaquine treatment

15 NSG mice with ectopic livers were infected with 3.3×10^5^ *P. falciparum* sporozoites (Seattle Children’s Research Institute, Seattle, WA), administered intravenously. Mice were grouped as follows to determine primaquine (PQ) prophylaxis efficiency through quantitative PCR (qPCR) analysis: *P. falciparum* only (n=3), *P. falciparum* + PQ 50 mg/kg for 3 days (n=3), and *P. falciparum* + PQ 30 mg/kg for 7 days (n=3). For quantitative reverse-transcription PCR (qRT-PCR) analysis of PQ prophylaxis efficiency the following groups were evaluated: *P. falciparum* only (n=3), and *P. falciparum* + PQ 30 mg/kg for 7 days (n=3). Dosing of PQ started one day before infection and continued until either day 3 or day 5 post sporozoite inoculation, as specified. All PQ used throughout these studies was synthesized by collaborators at Walter Reed Army Institute (WRAIR), reconstituted in PBS (HyClone, Marlborough, MA) and adjusted for salt concentration. Mice were sacrificed at designated time points with ectopic huLivers removed and tissue divided into <30 mg sections and placed in Allprotect Tissue Reagent (Qiagen, Hilden, Germany) for stabilization prior to extraction. Tissue sections were homogenized using 1 mm Zirconia/Silica beads (BioSpec Products, Bartlesville, OK) in a MagNA Lyser (Roche Life Science, Penzberg, Germany). For DNA extraction the QIAamp Kit was utilized (Qiagen, Hilden, Germany), following the manufacturer’s protocol, with quantity and quality determined using a NanoDrop spectrophotometer (Thermo Fisher, Waltham MA). qPCR was performed to quantify *P. falciparum* 18S genome copies with previously published primer and probe sequences[38]. The qPCR protocol was performed using iTAQ Universal Probes Supermix (Bio-Rad Laboratories, Hercules, CA) following the manufacture’s protocol. Parasite 18S genome copies were quantified using a standard curve with a plasmid control. For qRT-PCR analysis RNA was extracted using the RNeasy Plus Mini Kit (Qiagen, Hilden, Germany), following the manufacturer’s protocol. The isolated RNA was quantified using a NanoDrop spectrophotometer (Thermo Fisher, Waltham MA). RNA was reverse transcribed using the High-Capacity cDNA Reverse Transcription Kit (Applied Biosystems, Foster City, CA). The resulting cDNA was used for amplification using previously published *P. falciparum* primers and probe sequences [39], with the qRT-PCR protocol was performed using iTAQ Universal Probes Supermix (Bio-Rad Laboratories, Hercules, CA) with conditions following the manufacturer’s protocol. Parasite 18S copies were quantified using a standard curve with a plasmid control, which was generated from *P. falciparum* parasite DNA using a TOPO-TA cloning kit (Thermo Fisher, Waltham, MA) and previously published sequences [38]. All PCRs were performed on a an iCycler thermocycler equipped with an iCycler iQ real-time PCR detection system (Bio-Rad Laboratories, Hercules, CA).

#### *P. vivax* infections

*Ectopic liver infection with fresh P. vivax sporozoites:* 20 NSG mice with ectopic livers were infected with fresh *P. vivax* sporozoites at Mahidol University, Bangkok, Thailand. Sporozoites were generated by methods previously published [28]. Mice were infected with between 8.8×10^5^ and 1.08×10^6^ sporozoites administered i.v., with groups sacrificed on days 3 (n=5), 5 (n=4), 7 (n=5), and 9 (n=5). Ectopic liver tissue was sectioned and preserved for histology (snap frozen or formalin fixed) and sectioned into <30mg portions and placed in Allprotect Tissue Reagent (Qiagen, Hilden, Germany) for tissue stabilization prior to RNA extraction and subsequent qRT-PCR analysis.

##### P. vivax ectopic liver infection quantification through 18S qRT-PCR

Ectopic huLiver tissue was homogenized, RNA extracted and cDNA generated as previously described in text. cDNA was used for amplification with previously published *P. vivax* primers and probe sequences [40]. Parasite 18S copies were quantified using a standard curve with a plasmid control, which was generated from *P. vivax* parasite RNA using a TOPO-TA cloning kit (Thermo Fisher, Waltham, MA). The qPCR was performed using TaqMan Gene Expression Master Mix (Applied Biosystems, Foster City, CA) following the manufacturer’s protocol. All reactions were run using a ViiA 7 Real-Time PCR system (Applied Biosystems, Foster City, CA).

##### Immunofluorescence staining of UIS4

*P. vivax* infected ectopic liver tissue was formalin fixed, as previously described in text and sectioned. Tissues were incubated in blocking buffer (0.03% TritonX-100, 1% (w/v) BSA in 1 × PBS) with the mouse recombinant antibody anti-rUIS4 (1 μg ml−1, 1:10,000-fold dilution, kindly provided by Dr. Noah Sather, Seattle Children’s Research Institute, WA, USA) overnight at 4°C. Sections were washed thrice with 1× PBS then stained with the secondary antibody goat anti-mouse Alexa Fluor® 555 Plus conjugate (2 μg ml−1, 1:1000-fold dilution; Cat No. A32727, Molecular Probes, ThermoFisher, Waltham, MA, USA) overnight at 4°C. Samples were subsequently incubated for 1 h at room temperature using the nuclear stain Hoechst (10 μg ml−1, 1:1000-fold dilution) then washed thrice, and filled with 1× PBS for imaging and storage. Liver sections were imaged in bright field, TRITC, and DAPI channels at 10x or 20x magnification using the Operetta CLS high-content imaging system (Perkin Elmer, Waltham, MA, USA). Captured images were analyzed using Harmony software 4.9 (Perkin Elmer, Waltham, MA, USA) and scale bars were added using FIJI (ImageJ, version 1.53m).

#### Histology image acquisition and analysis

Unless otherwise noted, sections were visualized on an Eclipse TE 2000 microscope (Nikon, Tokyo, Japan). Images were collected using a wide field lens and NIS Elements version 4.2 software and viewed with NIS Elements Viewer version 4.11.0 software (all from Nikon). Fluorochrome settings were set to equivalent intensities among all samples for comparison and analysis. In addition, all files were exported using equivalent settings and file formats.

### Quantification and statistical analysis

All figures were generated using GraphPad Prism 8.0 software. Data is shown as mean ± SEM. For linear regression modeling, the coefficient of determination, R^2^, was defined using GraphPad Prism 6.0 for Mac.

Differences between route of infection, model (ectopic huLiver using HC-04 wild-type cell line, ectopic huLiver using HC-04 modified cell line, and FRG-huHep), and *P. falciparum* NF54HT parasite used (non-mouse adapted, mouse-adapted, and P_0_ NF54HT-GFP-luc) were analyzed with mixed effects models to account for repeated measurements in the mice. Prior to fitting the models, copy number was log_10_ transformed and all models were fit using SAS software (version 9.4; SAS Institute Inc., Cary, NC, USA). The final models included fixed terms for group, time, and the interaction between group and time with a random effect for mouse that used an unstructured covariance matrix. Groups and time points with only one mouse were excluded because they could not be reasonably used to test for differences between groups. To directly test differences between specific groups, hypothesis tests (*t-*tests) were conducted using contrast statements derived from the final models. Type III F-tests were used to evaluate overall differences. A *p-*value of < 0.05 was considered statistically significant.

## Results

### Ectopic human liver formation and characterization in NOD/SCID/ *IL2rg^null^* (NSG) mice

The HC-04 hepatocyte cell line was first evaluated for the ability to generate ectopic tumors in NSG mice. To do this 5.0×10^6^ cells were injected subcutaneously (s.c.) into NSG mice with tumor growth monitored over time using caliper measurements. Ectopic huLiver development was recorded with the first measurable huLiver appearing at 9 days post injection and followed through 22 days (Figure 1A). Next, we wanted to confirm vascularization of the tumor, which is a necessary element for sporozoite mobilization to the liver and required for malaria therapeutic delivery and absorption. Ectopic huLivers were shown to be vascularized with evidence of intratumoral microvessels as shown by laminin staining (Figure 1B).

**Figure 1.**
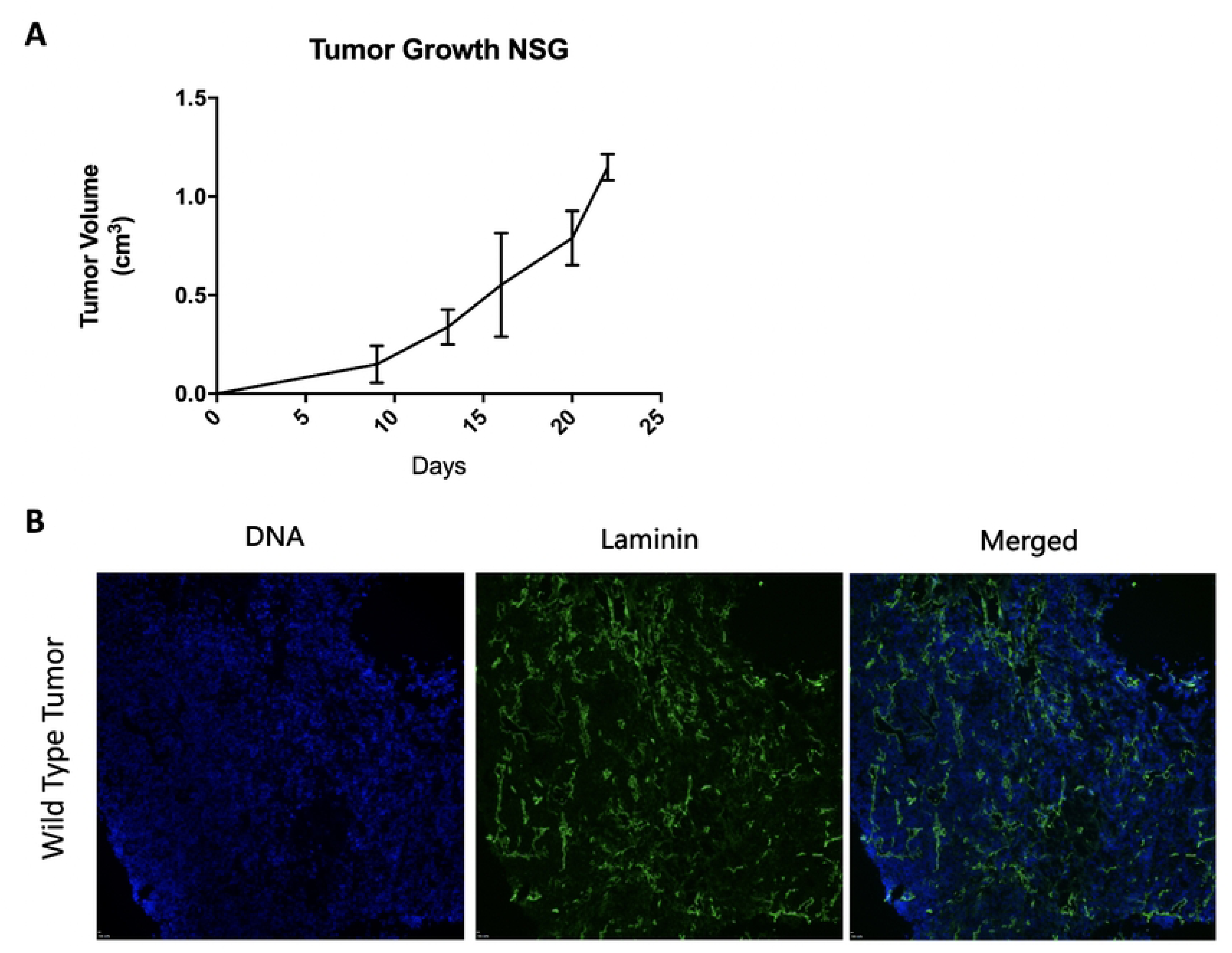
Characterization of the HC-04 cell line. **(A)** Ectopic huLiver (n=5) growth curve. Ectopic huLiver growth was followed for up to 22 days after s.c flank injection of 5.0×10^6^ HC-04 cells in NSG mice. **(B)** Vascularization of the ectopic huLiver by immunofluorescence labeling. DNA was visualized using DAPI (blue), and polyclonal rabbit anti-laminin Dylight 488 antibody (PA5-22901 Invitrogen) was used to show vascularization of the ectopic huLivers (green).

To further evaluate the vascularization of our ectopic huLiver model, LS infection with the genetically modified *P. berghei*-Luc sporozoites, a rodent *Plasmodium* species, that express the luciferase transgene [24] were used. *P. berghei* was selected over *P. falciparum* because liver development takes approximately 2 days as compared to 7 days. These *P. berghei*-Luc sporozoites have been genetically modified to express luciferase in a similar way as the *P. falciparum* NF54HT strain [41].

Ectopic huLiver mice were infected intratumorally (i.t.) with three different doses of *P. berghei*-Luc sporozoites. Luciferase activity in the liver was visualized through whole body imaging of the NSG mouse, using the *In Vivo* Imaging System (IVIS® Lumina LT *In Vivo* Imagining System, PerkinElmer) at 44 hours post-infection (hpi). Luciferase activity was detected in the ectopic huLiver of 1 out of 3 mice by IVIS (data not shown), which indicated that the ectopic huLiver is permeable to luciferin. Importantly, this IVIS result showed that sporozoites injected into the huLiver could travel through the bloodstream and infect the mouse liver, complementing the results of vascularization staining. *P. berghei*-Luc infection was additionally quantified by qRT-PCR to detect *P. berghei* 18S rRNA in both the ectopic huLiver and mouse liver. qRT-PCR results showed that *P. berghei* 18S rRNA was detectable in all infected mice, in both the ectopic huLivers and mouse livers (Supplementary Table 1).

These results show that qRT-PCR has a higher sensitivity than the IVIS luminescence read-out in our model, as luciferase activity was detected in only 1 out of 3 ectopic huLiver mice, suggesting that IVIS may not be an adequate method for determination of LS infection in this model.

Comparison of infection routes for NSG ectopic huLiver and FRG-huHep mice using the NF54HT-GFP-luc *P. falciparum* strain

To compare infection routes for *P. falciparum* we investigated if direct inoculation of sporozoites either through i.t injection of the ectopic huLiver mice or intravenous (i.v.) injection of FRG-huHep mice improved infection levels over a mosquito bite challenge, the natural route of infection. FRG-huHep mice acted as controls to the ectopic huLiver mice in these experiments.

First, NF54HT-Luc *P. falciparum* parasites were evaluated for the retention of luciferase expression. To do this an NSG mouse that had been engrafted with huRBCs to reach a 70% engraftment level, was injected with NF54HT-Luc *P. falciparum* infected blood, and imaged 8 minutes after luciferin was injected intraperitoneally (i.p). The IVIS imaging system, confirmed parasite luciferase activity (data not shown). Using NF54HT-GFP-luc *P. falciparum* sporozoites, three infection routes were compared: mosquito bite of ectopic huLiver mice (n=4) and FRG-huHep mice (n=3), injection of 100,000 freshly dissected sporozoites via i.t. (ectopic huLiver n=1) or i.v. (FRG-huHep n= 1).

A side-by-side comparison infection experiment was conducted. All mice were engrafted with huRBCs starting at day 5 post-infection, to reach engraftment of at least 70% by day 7, ready to be infected with the first wave of merozoites being released from the liver. At 6 days post-infection LS infection progression was assessed by IVIS (data not shown). Transition to blood-stage infection was monitored from 8 to 10 days post infection (dpi) using qRT-PCR to detect *P. falciparum* 18S rRNA in blood. Using IVIS, luciferase activity was detected above background for all FRG-huHep mice. However, none of the ectopic huLiver mice showed luciferase activity (data not shown). qRT-PCR to detect *P. falciparum* 18S rRNA in the blood showed that all ectopic huLiver and FRG-huHep mice, irrespective of the infection route had detectable 18S rRNA levels (Figure 2A). At 8 dpi, FRG-huHep mice had a higher infection with 10^11^ *P. falciparum* 18S copies/mL, as compared to ectopic huLiver mice with 10^10^ *P. falciparum* 18S copies/mL, though this difference was not statistically significant (Figure 2B). 18S RNA levels in the blood appeared to plateau by 9 dpi, staying stable to 10 dpi, with infection levels similar for all 3 groups at 10^10^ *P. falciparum* 18S copies/mL. Parasite levels obtained from sporozoite injection were similar to those obtained from the mosquito bite infection, demonstrating that sporozoites were infectious after dissection and injection (both i.t. and i.v.) (Figure 2A). Based on comparable infection levels we decided to use the mosquito bite as the route of infection in subsequent experiments. This eliminated the stress of dissection on the sporozoites, in addition to the time dependence required in obtaining viable sporozoites by dissection, thus simplifying the infection process. Transition to blood-stage was chosen as a readout for this model, as this was indicative of complete and successful LS development which culminated with the formation of merozoites that could infect huRBCs, as is observed during a natural infection process.

**Figure 2.**
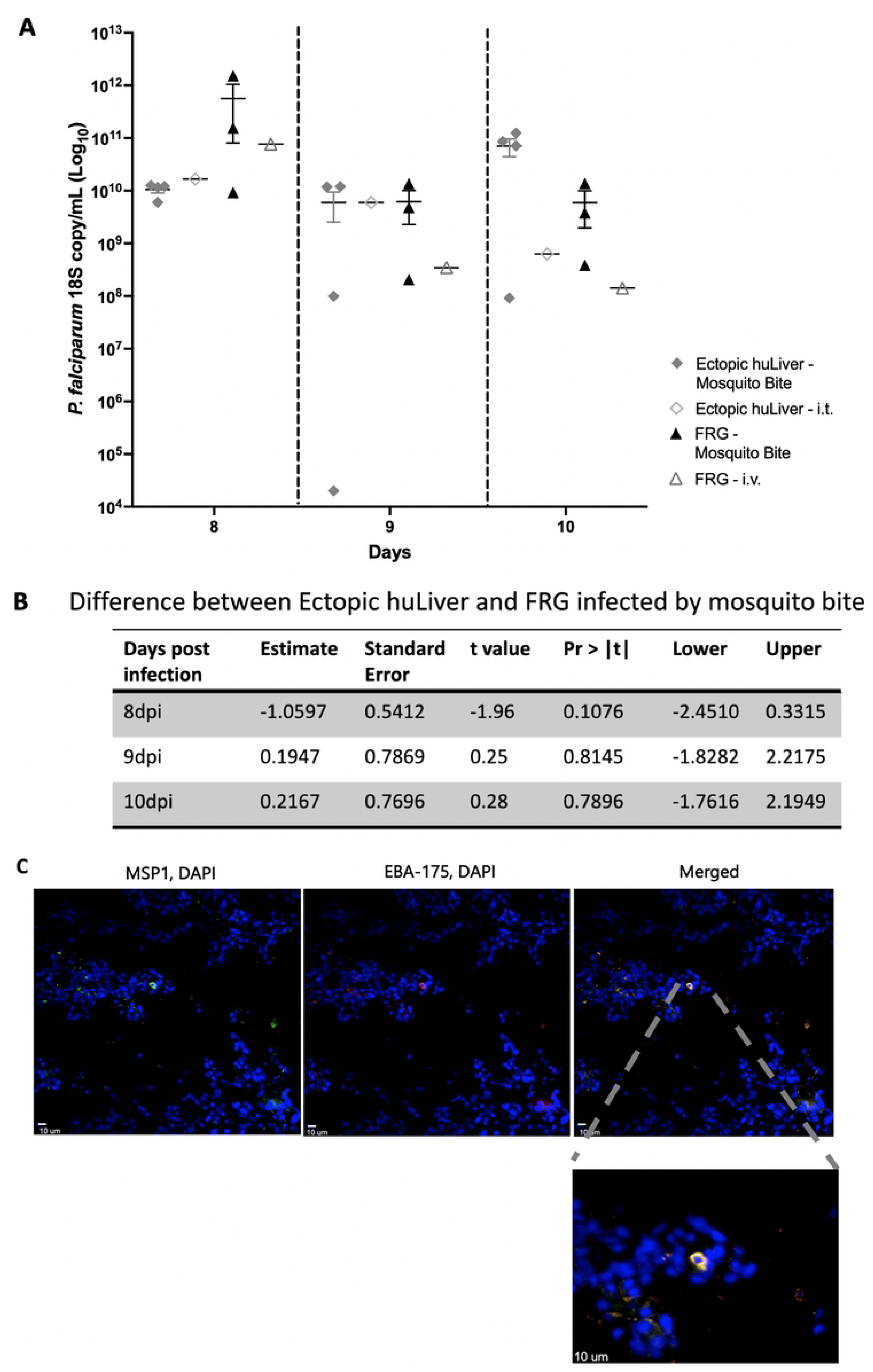
Route of infection comparison in NSG engrafted ectopic huLiver and FRG-huHep mice with the *P. falciparum* strain NF54HT-GFP-luc. Ectopic huLiver engrafted NSG, and FRG-huHep mice were infected with the *P. falciparum* strain NF54HT-GFP-luc. Routes of infection were compared between the ectopic huLiver and FRG-huHep mice. For comparison ectopic huLiver mice (n=4) and FRG-huHep mice (n=3) were infected via mosquito bite challenge and compared to infection from sporozoites (100,000 NF54HT-GFP-luc) administered via i.t (ectopic huLiver n=1) or i.v. (FRG-huHep n= 1). IHC and qRT-PCR analysis were performed to validate and characterize the infection. (A) Blood-stage infection was quantified using qRT-PCR to detect *P. falciparum* 18S rRNA transcripts in blood samples collected at 8, 9, and 10 dpi in both ectopic huLiver engrafted NSG and FRG-huHep mice. (B) Statistical analysis to demonstrate differences in infection levels between FRG-huHep and ectopic huLiver mice infected by mosquito bite at 8, 9, and 10. (C) Immunohistochemistry analysis using anti-MSP1 and anti-EBA-175 as markers for maturing merozoites, DAPI was used to stain cell DNA at 7 dpi.

Immunohistochemistry (IHC) was performed on flash frozen huLiver sections 7 days post infection with 1×10^7^ cryopreserved *P. falciparum* sporozoites for Merozoite Surface Protein 1 (MSP1) and Erythrocyte-Binding Antigen-175 (EBA-175). Late LS development is marked by the expression of MSP1 and EBA-175, which are both maturing merozoite markers [8]. The co-staining of both antibodies at 7 dpi indicated complete LS development had occurred, and DAPI staining showed a clear host hepatocyte nucleus. The positive IHC staining of MSP1 and EBA-175 is consistent with complete LS development in the ectopic huLiver model (Figure 2C).

### Characterizing ectopic liver metabolism and evaluating primaquine as an effective therapeutic for chemoprophylaxis in the ectopic huLiver model

To characterize the capability of this model to successfully uptake anti-malarial compounds, we looked at expression of CYP 2D6 through RT-PCR analysis and hepatic asialoglycoprotein receptor 1 (ASGPR1) expression through IHC in the HC-04 cell line. Additionally, the pharmacokinetics (PK) of primaquine (PQ) metabolism in the mouse liver was compared to the ectopic liver. As a follow-up to these experiments, we tested the ability of primaquine to function prophylactically in this model against *P. falciparum* infection.

IHC staining for ASGPR1 on the ectopic liver hepatocytes showed positive expression of the transmembrane C-type lectin receptor (Figure 3A) that has been extensively used for targeted delivery of therapeutics to the liver [42]. Expression levels of CYP 2D6, a gene necessary for the metabolism and generation of the active metabolite for primaquine and directly linked to efficacy [43], was measured in the HC-04 cell line and compared to commercially available pooled human liver RNA. It was found that CYP 2D6 is expressed in the HC-04 cell line, although at a 200-fold decrease to pooled human liver RNA (Figure 3B).

**Figure 3.**
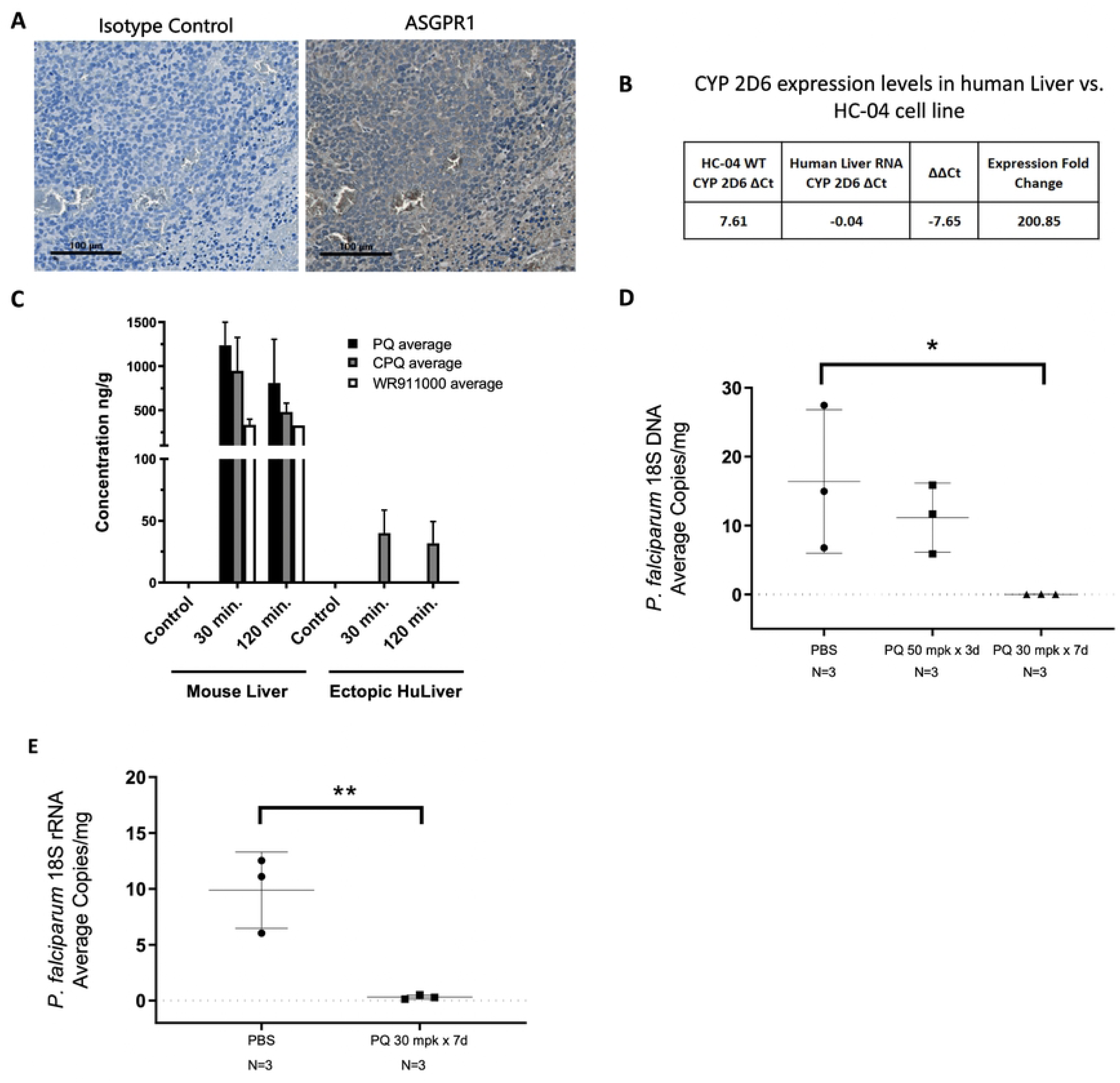
Pharmacokinetics of primaquine in the ectopic huLiver model and evaluation of primaquine as a prophylactic for *P. falciparum* infection. (A) Immunohistochemical (IHC) staining for anti-ASGPR1 in ectopic huLiver tissue (Rb anti-ASGPR1 1:500) compared to an isotype control (RB IgG 1:500). (B) Expression of CYP 2D6 in the HC-04 cell line compared to pooled human liver RNA. (C) Pharmacokinetic comparison of primaquine in ectopic huLiver and mouse liver tissue harvested at 30 and 120 minutes following a single dose of PQ at 30 mpk (p.o.) (D) *P. falciparum* 18S DNA copies per mg of ectopic huLiver tissue (*p-value:* 0.0415). (E) *P. falciparum* 18S rRNA copies per mg of ectopic huLiver tissue (*p-value:* 0.0084).

The PK profile of PQ in the ectopic liver is important as PQ acts on parasites within the liver and therefore must be able to penetrate the site of infection. In this study, ectopic huLiver mice were dosed with either PBS (control) or PQ at 30 milligrams per kilogram (mpk) orally (p.o.) and sacrificed at 30 minutes and 120 minutes post dosing with both the ectopic huLiver and mouse liver tissue analyzed for the PQ metabolites: carboxy-PQ (CPQ) and the PQ orthoquinone WR911000. In treated mouse liver tissue, PQ metabolites remained high at both 30- and 120-minutes post dosing, with only CPQ detected in the ectopic liver at both timepoints. These results suggest that metabolism is occurring via the mouse liver, and the CPQ is able to penetrate the ectopic huLiver tissue (Figure 3C).

To assess whether parasites were actively replicating in the huLiver, RNA was extracted from the ectopic livers and qRT-PCR to analyze *P. falciparum* 18S rRNA transcripts in PQ treated vs. untreated groups was performed. Mice were prophylactically treated with PQ at 30 mpk for 7 days (n=3) starting one day prior to cryopreserved *P. falciparum* GFP-luc sporozoite inoculation, alongside a vehicle control (PBS, n=3) group. All mice were euthanized at 7 days post infection with ectopic livers harvested for DNA and RNA extraction. Results from the qPCR showed a statistically significant decline in transcripts in the PQ at 30 mpk for 7 days treatment group compared to the vehicle control group (*p-value:* 0.0415) (Figure 3D). qRT-PCR analysis of the PQ treated group compared to the vehicle control group was also found to be statistically significant (*p-value:* 0.0084) (Figure 3E). These results demonstrated that PQ acted as a causal prophylactic in the ectopic huLiver mice infected with *P. falciparum*.

### Receptor modified HC-04 cell line for ectopic-huLiver formation

While *P. falciparum* infection was observed in both the ectopic huLiver model and FRG-huHep mice, the infection levels were low. In order to address this, the possible effect of *P. falciparum* receptor levels on HC-04 cell line on infectivity was investigated. Studies have shown that the HC-04 cell line has low levels of CD-81 and EphA2 receptor expression, both of which are critical for *P. falciparum* infection [44–48]. To determine if increasing these receptor levels would improve infection rates in our ectopic huLiver model, we generated a HC-04 cell line with higher expression of both EphA2 and CD81; HC-04 EphA2^High^ CD81 ^High^.

The ability of the EphA2^High^ CD81 ^High^ modified and HC-04 wild-type (WT) cell lines to form ectopic huLivers in NSG mice was compared by injecting s.c. 5.0×10^6^ cells from each cell line and monitoring tumor growth over time.

The HC-04 EphA2^High^ CD81 ^High^ modified cell line created an ectopic huLiver in NSG mice that had similar growth dynamics, size (Supplementary Figure 1A), vascularization (Supplementary Figure 1B), and time to develop, as the WT HC-04 cell line. Additionally, immunofluorescent imaging (Supplementary Figure 1B), and qRT-PCR (Supplementary Figure 1C) confirmed that the ectopic liver formed using the HC-04 EphA2^High^ CD81 ^High^ cell line retained high receptor expression *in vivo*.

### Comparison of LS development and blood-stage transition between NSG ectopic huLiver infection and FRG-huHep infection with the NF54HT *P. falciparum* strain

The NF54HT *P. falciparum* strain infectivity was compared across three forms: (i) a non-mouse adapted NF54HT *P. falciparum* strain which did not express GFP or luciferase, (ii) a mouse-adapted NF54HT *P. falciparum* strain that had been previously passaged through mice for 10 cycles and had been shown to grow *in vivo* in the blood of humanized mice [33], and (iii) P_0_ from the genetically modified NF54HT-GFP-luc *P. falciparum* strain, which had one replication cycle completed in FRG-huHep. The P_0_ from the genetically modified NF54HT-GFP-luc *P. falciparum* strain was generated to test if one passage through mice improved the parasite infection fitness, as had been previously reported for blood-stage [33].

Infectability of ectopic huLivers generated with either the HC-04 wild-type cell line or the modified HC-04 EphA2^High^ CD81 ^High^ cell line was assessed and directly compared against FRG-huHep mice, which were used as positive controls. All infections were done by mosquito bite. In addition, to evaluate blood-stage transition, all the mice were engrafted daily with huRBCs to reach engraftment of at least 70% by day 7. Transition to blood-stage infection was determined starting at 8 dpi and followed over time using qRT-PCR to detect of *P. falciparum* 18S rRNA in blood. Experimental end-points for the ectopic huLiver mice were based on the size of the tumor (1.5 cm^3^, Supplementary Figure 1A) and ranged from 10-16 dpi.

Using qRT-PCR to detect *P. falciparum* 18S rRNA in blood, we first evaluated whether the non-mouse adapted NF54HT *P. falciparum* strain could transition from liver to blood-stage infection. Parasite levels in blood followed the expected replication kinetics, with *P. falciparum* 18S copies/mL around 10^9^ - 10^10^ from 8 to 12 dpi (Figure 4A). The mouse-adapted NF54HT *P. falciparum* strain was used to determine if passaging the parasite through mice would increase the overall infectivity, as previously reported for blood-stage [33]. The levels of parasite detected for this mouse-adapted strain were similar to the non-mouse adapted NF54HT *P. falciparum* strain, with approximately 10^8^ *P. falciparum* 18S copies/mL detected at 8 dpi which gradually increased, reaching a peak close to 10^9^ *P. falciparum* 18S copies/mL by 12 dpi (Figure 4B). These results indicated that in both the FRG-huHep and ectopic huLiver, the mouse-adapted *P. falciparum* NF54HT strain did not increase the overall infection levels.

**Figure 4.**
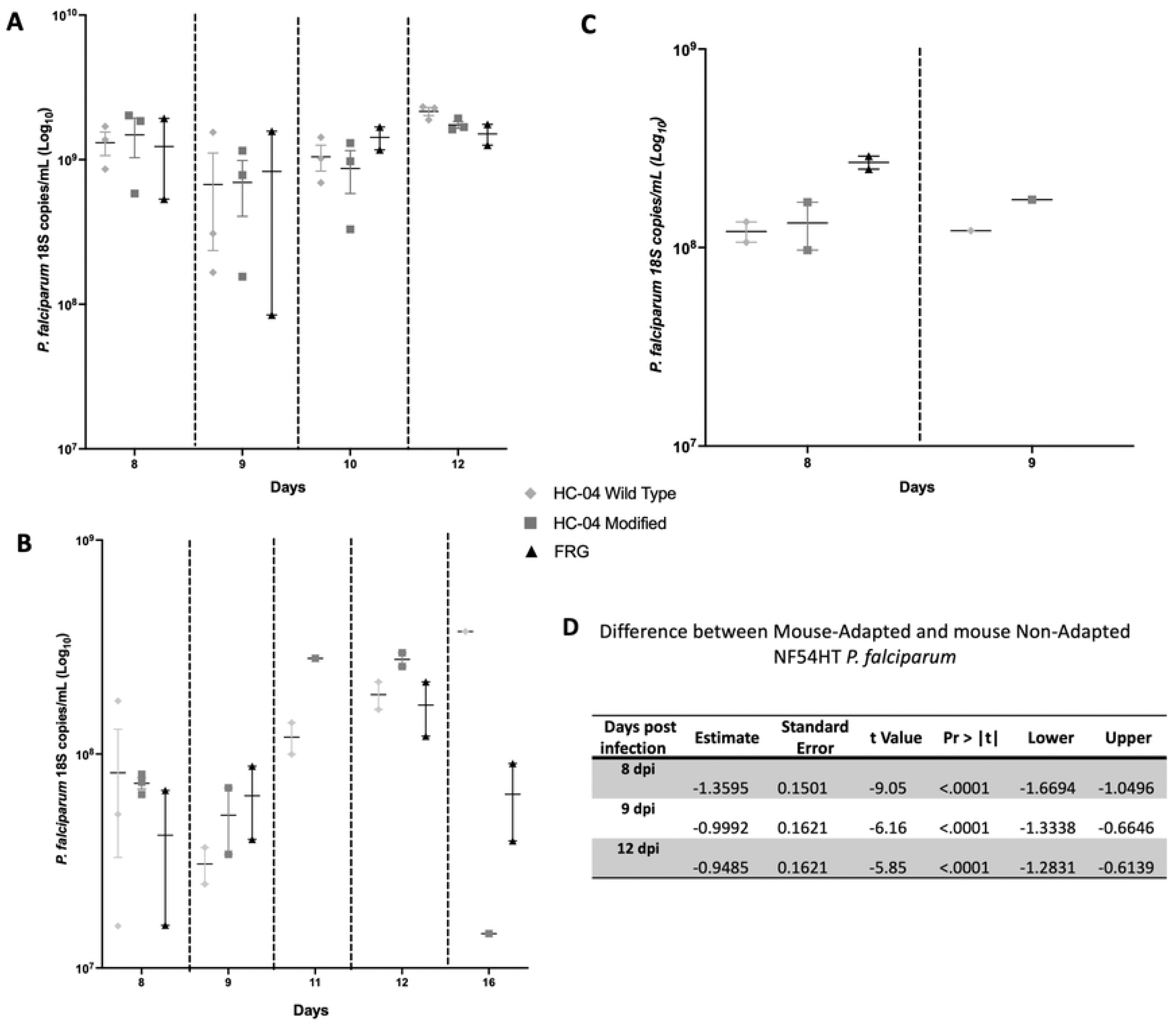
Comparison of NSG ectopic huLiver infection to the FRG-huHep infection model with the *P. falciparum* strain NF54HT with three adaptations. Transition from liver to blood-stage infection was quantified using qRT-PCR to detect *P. falciparum* 18S rRNA expression in both the ectopic huLiver engrafted NSG and FRG-huHep infection models. (A) *P. falciparum* 18S rRNA expression in mosquito bite infected mice from each model (HC-04 WT n = 3, HC-04 modified n = 3 FRG-huHep n =2) with the non-mouse adapted NF54HT *P. falciparum* strain. Blood samples were collected at 8 through 12 dpi. (B) *P. falciparum* 18S rRNA expression in mosquito bite infected mice from each model (HC-04 WT n = 3, HC-04 modified n = 3 FRG-huHep n =2) with the mouse adapted NF54HT *P. falciparum* strain. Blood samples were collected at 8 through 16 dpi. (C) Statistical Analysis showing differences between mouse-adapted and non-mouse adapted NF54HT *P. falciparum* at day 8, 9 and 12 post infection. (D) Ectopic huLiver engrafted NSG (HC-04 WT n = 2, HC-04 modified n = 2) and FRG-huHep mice (n =3) were infected by mosquito bite using P_0_ of NF54HT-GFP-luc, generated from the first completed life cycle passage within FRG-huHep. Blood samples were collected 8 and 9 dpi for qRT-PCR analysis of *P. falciparum* 18S rRNA expression

Finally, we used the P_0_ NF54HT-GFP-luc *P. falciparum* strain to infect both the FRG-huHep and ectopic huLiver mice. At 6 dpi, liver-stage infection was assessed using IVIS imaging, with luciferase activity detectable in two out of three FRG-huHep; however, no luminescence was observed for the ectopic huLiver mice (data not shown). Transition to blood-stage infection was followed over time in all mice using qRT-PCR to detect *P. falciparum* 18S rRNA. Results indicated all mice had infection, with ectopic huLiver albeit at a slightly lower infection than FRG-huHep mice (Figure 4C). Infection levels were between 10^8^ parasites for ectopic huLiver and 10^9^ parasites for FRG-huHep mice. These results demonstrated that one completed life cycle in the FRG-huHep mice was not enough to increase the infection efficiency. It remains to be investigated if additional parasite passages through the mice would increase infectivity.

Overall, the non-mouse adapted NF54HT *P. falciparum* strain was found to be significantly better (p-value: 0.0218) than the mouse-adapted NF54HT *P. falciparum* and the NF54HT-GFP-luc *P. falciparum* P_0_ (Figure 4D). Additionally, all the infectivity differences observed between the ectopic huLiver model, with either the HC-04 wild-type cell line or the modified HC-04 EphA2^High^ CD81 ^High^ cell line, and the FRG-huHep model were not statistically significant. Indicating that the ectopic huLiver model had infection levels that were similar to those seen in the FRG-huHep mice and that the increase in CD81 and EphA2 receptor expression in the HC-04 cell line did not improve infection efficiency.

### *Plasmodium vivax* infection of ectopic huLiver mice using fresh sporozoites

Lastly, we wanted to determine if the ectopic huLiver model could also be utilized to study *P. vivax* infection. Fresh wild-type sporozoites were obtained from dissected mosquito salivary glands of mosquitos which had taken a blood meal from *P. vivax* infected blood. Mice were infected i.v. with between 8.8 × 10^5^ and 1 × 10^6^ sporozoites and then divided into 4 groups with mice euthanized and ectopic huLiver tissue harvested on days 3, 5, 7, and 9 post infection. Ectopic huLiver tissue was sectioned for immunohistochemistry and RNA extraction for qRT-PCR to detect *P. vivax* 18S rRNA. The results from the qRT-PCR showed parasite transcripts increased from day 3 and 7 and then dropped off precipitously on day 9 (Figure 5A). The observed pattern may reflect an expansion of LS parasites, which are increasing in copy number prior to merozoite release from hepatocytes. On day 9 it is likely that the only remaining parasites are dormant hypnozoites, which have less transcriptional activity. To look for evidence of hypnozoites, 7 days post infection tissue was sectioned and stained for rUIS4, a parasitophorous vacuole membrane (PVM) stain specific for *P. vivax*, which can allow visual differentiation of LS schizonts and hypnozoites. Hypnozoites were detected as evidenced by staining with the rUIS4 antibody (Figure 5B). These experiments demonstrate that the ectopic huLiver model supported *P. vivax* infection including hypnozoite formation.

**Figure 5.**
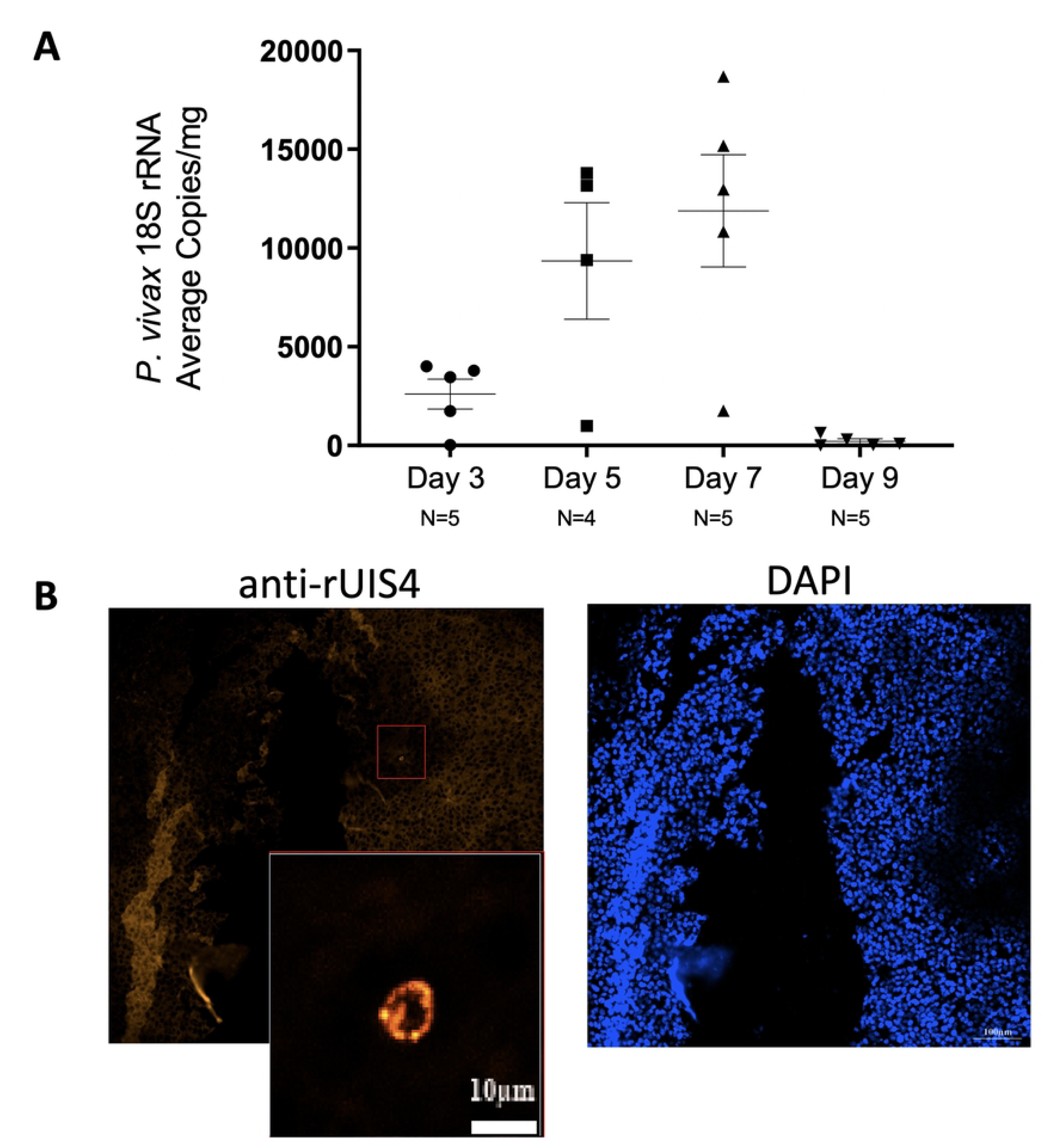
Ectopic huLiver mice infected with freshly dissected *P. vivax* sporozoites. NSG ectopic huLiver mice were i.v. infected with between 8.8 × 10^5^ and 1 × 10^6^ *P. vivax* sporozoites. Mice were euthanized on day 3, 5, 7 or 9 post infection with ectopic liver tissue harvested for IHC and RNA extraction to determine 18s rRNA transcripts levels by qRT-PCR. **(A)** Quantification of parasite ectopic huLiver burden by qRT-PCR of *P. vivax* 18S rRNA. **(B)** Immunohistochemistry analysis at 7 days post infection using anti-rUIS4 and DAPI at 20X magnification. White scale bar represents 100 µm and grey scale bar represents 10 µm.

## Discussion

Malaria presents a major global health burden causing roughly half a million deaths per year and placing approximately half the world’s population at risk of infection. One of many challenges associated with reducing cases worldwide is the lack of adequate models available to study human-specific *Plasmodium* infection. Existing preclinical animal models for examining the liver stage phase of human malaria have significant limitations that hinder the investigation of parasite biology and drugs that target the liver stage as well as the hypnozoite forms responsible for relapsing malaria. Several recent studies describing humanized mouse models have demonstrated that human *Plasmodium* infection can occur in a humanized liver [8,13,15,49,50]. However, these models utilize primary human hepatocytes, which introduce donor-to-donor variability, yielding reproducibility issues. They also require surgery or other considerable technical manipulation which makes them challenging for use in antimalarial drug discovery. Presented here is a novel model for both *P. falciparum* and *P. vivax* LS infection-, the ectopic huLiver mouse model. This model is a superior addition to currently available *in vivo* research models as it uses a human hepatocyte cell line that can be infected by both *P. falciparum* and *P. vivax,* is moderate-cost, easy-to-manipulate, reproducible, supports testing of anti-malarial therapeutics, and could serve to help further elucidate LS parasite biology.

The ectopic huLiver mouse model is a tumor model in which tumors are generated using the HC-04 cell line [11]. These cells were first described as immortalized, yet when engrafted subcutaneously in immunodeficient NSG mice, these cells were observed to divide with unrestricted growth, and developed a vascularized blood vessel network characteristic of tumor formation more indicative of a transformed cell line.

One key aspect of this model is vascularization as it supports tumor growth, allows for sporozoites and antimalarial compounds and metabolites to permeate the site of infection, and facilitates the transition from LS to blood-stage as merozoites must egress from the ectopic huLiver into the bloodstream. The ability of the model to facilitate the distribution of experimental compounds to the site of infection was confirmed by pharmacokinetic analysis of PQ metabolism in the ectopic huLiver tissue. This was again confirmed by experiments where PQ successfully acted as a causal prophylactic in ectopic huLiver mice infected with *P. falciparum*, indicating that this model could be useful in the assessment of anti-malarial therapeutic efficacy. Another feature of this model is the presence of the liver enzyme metabolizer CYP 2D6. It has been shown that the cytochrome P450 (CYP) hepatic enzyme CYP 2D6 is required for the metabolism of PQ and potentially other 8-aminoquinolines, excluding tafenoquine [51, 52]. In models using repopulation with primary hepatocytes variability in CYP 2D6 function is a result of donor dependent levels, making standardization of such models challenging for evaluating anti-malarials consistently across experiments. Additionally, IHC staining of the HC-04 cell line confirmed the presence of the receptor asialoglycoprotein receptor 1 (ASGPR1), an essential receptor for hepatocyte targeted drug delivery, which is associated with drug uptake in cancer cells [53]. While not previously studied in the context of malaria LS treatment, ASGPR1 has been extensively utilized as a target for human hepatocellular carcinoma therapies [36].

The best characterized humanized mouse model for LS malaria infection is the FAH*^−/-^*, FAH*^−/-^*, Rag2*^−/-^*, IL2rγ*^−/-^* (FRG) immunodeficient mouse which supports both P. *falciparum* and *P. vivax* LS and blood stage transition [8]. This FRG mouse model has been shown to support infection from both mosquito bite and i.v. sporozoite injection [8, 30].

Ectopic huLiver infection by both mosquito bite and intravenous injection was successful. Nevertheless, infection by mosquito bite was selected when the option was available as it is the natural route of infection, and using this method not only simplified the process by eliminating the time-consuming manual labor required for preparation of sporozoites but also imposed less stress on the system. One of the challenges associated with using the mosquito bite method of introducing infection in an *in vivo* model for drug testing is the issue of dose reproducibility, as the amount of sporozoites that each mouse receives cannot be controlled. However, a recent study has shown that mosquito bite challenge can be optimized. Using the FRG huHep model, mice were challenged with 40 infected mosquitos 10 minutes mice which resulted in consistent liver stage burdens [30], thus optimizing a method that could be applicable in the ectopic huLiver model in future studies.

In optimizing a method for quantifying infection, the IVIS was found to lack the sensitivity necessary for detection of NF54HT-GFP-luc *P. falciparum* infected hepatocytes in the ectopic huLiver model. This was unexpected as previously published papers using the FRG mouse model as well as our own data have consistently shown IVIS detection in NF54HT-GFP-luc *P. falciparum* LS infection [30,54,55]. The ectopic huLivers were vascularized and able to support detection of *P. berghei-Luc* by IVIS imaging, suggesting that the lack of detection of an IVIS signal was not due to lack of permeabilization of the tumor tissue (data not shown). However, despite the lack of detectable luciferase activity, qRT-PCR for detection *Plasmodium* 18S RNA confirmed the presence of infection with both *P. falciparum* and *P. vivax* in ectopic huLiver tissue. Transcripts for *Plasmodium* 18S RNA were also detected in the of blood of *P. falciparum* infected ectopic huLiver mice, but remains to be investigated with *P. vivax*. It is therefore possible that the levels of infection were below the levels of detection for IVIS imaging. Furthermore, we also investigated the efficiency of flow cytometry to measure *P. falciparum* transition to blood stage infection, this technology also proved to have very low sensitivity even for FRG-huHep mice (data not shown). Despite these limitations, detection of *Plasmodium* by tracking the liver to blood stage transition using qRT-PCR to detect 18SRNA in blood would provide: (i) evidence that the LS phase was completed and successfully produced merozoites capable of infecting huRBCs (ii) infection amplification during blood stage; an infected huRBCs (ihuRBCs) produces infectious merozoites that go on and infected naïve huRBCs augmenting infection, and (iii) it allows the opportunity to follow infection over time; only a small sample of blood must be collected at each time point and mice can be maintained for a longer duration.

Two variations of the HC-04 cell line for *P. falciparum* were compared for infection efficiency; the wild-type HC-04, and the modified HC-04 which was engineered to express elevated levels of both CD81 and EphA2 receptors. The experiments described here suggest that neither of these receptors increased the overall infection efficiency of the model. It has recently been reported that CD81 creates microdomains that are either beneficial for the parasite infection [47, 48] or regulating another unidentified receptor [45, 46], these studies could explain why artificially increasing the receptor had no effect on infection levels in our model. Another possibility is that there is an initial uptick in sporozoite invasion in huHep as CD81 has been shown to be linked invasion, but that the parasite does not have enough cofactors present to develop or survive beyond the earliest stages. Conversely, EphA2 has been shown to directly engage with *P. falciparum* 6-cys fold proteins [44]. One hypothesis as to why elevated EphA2 had no effect on infection efficiency is that *P. falciparum* replication *in vivo* requires a more stringent microenvironment that the receptor alone is not able to overcome, although this remains to be elucidated.

*P. falciparum* experiments were conducted using the NF54HT strain in both the mouse-adapted and non-mouse adapted form as well as the NF54HT-GFP-luc strain. We were interested in investigating the infection efficiency of the NF54HT-GFP-luc *P. falciparum* strain compared to the non-mouse adapted NF54HT strain. The data presented here exhibited a similar trend, where the non-mouse adapted NF54HT *P. falciparum* strain had statistically significant increased infection levels compared to the NF54HT-GFP-luc and the mouse adapted strain (*p-value:* 0.0218). Previous studies have reported similar results when comparing mouse adapted and clinical isolates of *P. falciparum*, indicating that the parasite does not require adaptation to a mouse host to achieve a robust infection [56].

The *Plasmodium vivax* experiments were conducted utilizing wild-type parasites sporozoites freshly dissected from salivary glands of mosquitos which had fed on blood from an infected individual. Using this clinically isolated strain of *P. vivax,* we found enhanced LS infection efficiency in the ectopic huLiver tissue as compared with infections resulting from cryopreserved *P. falciparum* lab-adapted strains. Therefore, we hypothesize that the use of clinically isolated *Plasmodium* strains could have greater infection efficiency than the laboratory-adapted strains investigated in this study. Additionally, pairing wild-type isolates of *P. falciparum* and *P. vivax* with mosquito bite transmission in this model could prove to be a promising avenue for testing anti-malarial compounds against more diverse and clinically relevant strains.

The main limitations of our experiments with both *P. falciparum* and *P. vivax* were the small sample size of infection groups and the lack of consistency pertaining to parasite sourcing. A notable bottleneck in *Plasmodium* LS research is that few laboratories have the ability to maintain a mosquito vivarium due to financial or geographical constraints and often access is limited due to high demand where they do exist. It is for this reason that despite our preference to elicit infection using the mosquito bite method, some of the experiments described in this text were conducted with dissected cryopreserved sporozoites, notably the prophylactic primaquine experiments with *P. falciparum*. Additionally, the observed size of the liver-stage schizonts in the ectopic huLiver model at 7 days post infection with *P. falciparum* had a diameter of 10μm, compared to previously published data which have shown 15-20μm diameter [57]. Another limitation in regards to studying *P. vivax* infection in the ectopic huLivers is the time constraints associated with using a tumor model due to tumor growth dynamics to recapitulate LS infection, as this would restrict the ability to study hypnozoite biology. Hypnozoites, while thought to form at the initiation of infection and distinguishable from schizonts on day 3 by immunofluorescence staining would require treatment and time for clearance to be monitored [15]. Therefore, the ectopic huLiver model would ideally need to support infection for at least 14 days post infection. Future experiments are planned to transfer infected ectopic livers to new recipient NSG mice to sustain the infection.

Evidence for the strength of our ectopic huLiver model and its utility in the study of *P. falciparum* LS biology, liver to blood-stage transition, and as a drug discovery tool is based on our ability to measure ihuRBCs in mice. ihuRBCs are only detectable after merozoites invade and replicate inside huRBCs, making it a biologically relevant model to study *P. falciparum* LS infection. The use of the HC-04 cell line as the basis for the ectopic huLiver model intrinsically provides hepatic receptors required for drug uptake, allows for ease of use, reproducibility, and the potential to scale up for sizable therapeutic drug studies.

Briefly, we showed that the model is robust and biologically relevant and could be useful for drug development, both vaccine development and monoclonal antibody production, by providing a platform for therapeutic discovery pathways against *P. falciparum* infection and/or as a model for analysis of drug efficacy against LS or LS to blood-stage transition. The development of the ectopic huLiver model provides an additional tool in the anti-malaria armamentarium against human *Plasmodium* LS.

## Disclaimers

Material has been reviewed by the Walter Reed Army Institute of Research. There is no objection to its presentation and/or publication. The opinions or assertions contained herein are the private views of the author, and are not to be construed as official, or as reflecting true views of the Department of the Army or the Department of Defense.

Research was conducted under an approved animal use protocol in an AAALAC International-accredited facility in compliance with the Animal Welfare Act and all other federal statutes and regulations relating to animals and experiments involving animals, and adheres to principles stated in the Guide for Care and Use of Laboratory Animals, NRC Publication, 2011 edition.

## References

1. World Health Organization. World malaria report 2020: 20 years of global progress and challenges. Geneva; 2020 Licence: CC BY-NC-SA.0 IGO p. 299.

2. Vaughan AM, Kappe SHI. Malaria Parasite Liver Infection and Exoerythrocytic Biology. Cold Spring Harb Perspect Med. 2017 Jun;7(6):a025486.

3. Howes RE, Battle KE, Mendis KN, Smith DL, Cibulskis RE, Baird JK, Hay SI. Global Epidemiology of *Plasmodium vivax*. Am J Trop Med Hyg. 2016 Dec 28;95(6 Suppl):15–34.

4. Amino R, Giovannini D, Thiberge S, Gueirard P, Boisson B, Dubremetz JF, Prévost MC, Ishino T, Yuda M, Ménard R. Host Cell Traversal Is Important for Progression of the Malaria Parasite through the Dermis to the Liver. Cell Host & Microbe. 2008 Feb;3(2):88–96.

5. Garcia JE, Puentes A, Patarroyo ME. Developmental Biology of Sporozoite-Host Interactions in Plasmodium falciparum Malaria: Implications for Vaccine Design. Clinical Microbiology Reviews. 2006 Oct 1;19(4):686–707.

6. Zuck M, Austin LS, Danziger SA, Aitchison JD, Kaushansky A. The Promise of Systems Biology Approaches for Revealing Host Pathogen Interactions in Malaria. Front Microbiol. 2017 Nov 16;8:2183.

7. Adams JH, Mueller I. The Biology of *Plasmodium vivax*. Cold Spring Harb Perspect Med. 2017 Sep;7(9):a025585.

8. Vaughan AM, Mikolajczak SA, Wilson EM, Grompe M, Kaushansky A, Camargo N, Bial J, Ploss A, Kappe SHI. Complete Plasmodium falciparum liver-stage development in liver-chimeric mice. J Clin Invest. 2012 Oct 1;122(10):3618–28.

9. Mikolajczak SA, Sacci Jr JB, De La Vega P, Camargo N, VanBuskirk K, Krzych U, Cao J, Jacobs-Lorena M, Cowman AF, Kappe SHI. Disruption of the Plasmodium falciparum liver-stage antigen-1 locus causes a differentiation defect in late liver-stage parasites: Deletion of P. falciparum LSA-1 gene. Cellular Microbiology. 2011 Aug;13(8):1250–60.

10. Vaughan AM, O’Neill MT, Tarun AS, Camargo N, Phuong TM, Aly ASI, Cowman AF, Kappe SHI. Type II fatty acid synthesis is essential only for malaria parasite late liver stage development. Cellular Microbiology. 2009 Mar;11(3):506–20.

11. Brewer TG, Leelaudomlipi S, Coleman RE, Cui L, Jenwithisuk R, Yimamnuaychoke N, Rasameesoraj M, Udomsangpetch R, Sattabongkot J. Establishment of a Human Hepatocyte Line that Supports in vitro Development of the Exo-erythrocyticStages of the Malaria Parasites Plasmodium falciparum and P. vivax. The American Journal of Tropical Medicine and Hygiene. 2006 May 1;74(5):708–15.

12. Dumoulin PC, Trop SA, Ma J, Zhang H, Sherman MA, Levitskaya J. Flow Cytometry Based Detection and Isolation of Plasmodium falciparum Liver Stages In Vitro. Silvie O, editor. PLoS ONE. 2015 Jun 12;10(6):e0129623.

13. Sacci JB, Alam U, Douglas D, Lewis J, Tyrrell DLJ, Azad AF, Kneteman NM. Plasmodium falciparum infection and exoerythrocytic development in mice with chimeric human livers. International Journal for Parasitology. 2006 Mar;36(3):353–60.

14. Vaughan AM, Pinapati RS, Cheeseman IH, Camargo N, Fishbaugher M, Checkley LA, Nair S, Hutyra CA, Nosten FH, Anderson TJC, Ferdig MT, Kappe SHI. Plasmodium falciparum genetic crosses in a humanized mouse model. Nat Methods. 2015 Jul;12(7):631–3.

15. Mikolajczak SA, Vaughan AM, Kangwanrangsan N, Roobsoong W, Fishbaugher M, Yimamnuaychok N, Rezakhani N, Lakshmanan V, Singh N, Kaushansky A, Camargo N, Baldwin M, Lindner SE, Adams JH, Sattabongkot J, Kappe SHI. Plasmodium vivax Liver Stage Development and Hypnozoite Persistence in Human Liver-Chimeric Mice. Cell Host & Microbe. 2015 Apr;17(4):526–35.

16. Schäfer C, Roobsoong W, Kangwanrangsan N, Bardelli M, Rawlinson TA, Dambrauskas N, Trakhimets O, Parthiban C, Goswami D, Reynolds LM, Kennedy SY, Flannery EL, Murphy SC, Sather DN, Draper SJ, Sattabongkot J, Mikolajczak SA, Kappe SHI. A Humanized Mouse Model for Plasmodium vivax to Test Interventions that Block Liver Stage to Blood Stage Transition and Blood Stage Infection. iScience. 2020 Aug;23(8):101381.

17. Strom SC, Davila J, Grompe M. Chimeric Mice with Humanized Liver: Tools for the Study of Drug Metabolism, Excretion, and Toxicity. In: Maurel P, editor. Hepatocytes [Internet]. Totowa, NJ: Humana Press; 2010 [cited 2020 Jul 7]. p. 491–509. (Methods in Molecular Biology; vol. 640). Available from: http://link.springer.com/10.1007/978-1-60761-688-7_27

18. Bertilsson L, Dahl ML, Dalén P, Al-Shurbaji A. Molecular genetics of CYP2D6: Clinical relevance with focus on psychotropic drugs: *Molecular genetics of CYP2D6*. British Journal of Clinical Pharmacology. 2002 Feb;53(2):111–22.

19. Bissig KD, Han W, Barzi M, Kovalchuk N, Ding L, Fan X, Pankowicz FP, Zhang QY, Ding X. P450-Humanized and Human Liver Chimeric Mouse Models for Studying Xenobiotic Metabolism and Toxicity. Drug Metab Dispos. 2018 Nov;46(11):1734–44.

20. Katoh M, Sawada T, Soeno Y, Nakajima M, Tateno C, Yoshizato K, Yokoi T. In vivo drug metabolism model for human cytochrome P450 enzyme using chimeric mice with humanized liver. Journal of Pharmaceutical Sciences. 2007 Feb;96(2):428–37.

21. Potter BMJ, Xie LH, Vuong C, Zhang J, Zhang P, Duan D, Luong TLT. Differential CYP 2D6 Metabolism Alters Primaquine Pharmacokinetics. Antimicrobial Agents and Chemotherapy. 2015;59(4):8.

22. Azuma H, Paulk N, Ranade A, Dorrell C, Al-Dhalimy M, Ellis E, Strom S, Kay MA, Finegold M, Grompe M. Robust expansion of human hepatocytes in Fah−/−/Rag2−/−/Il2rg−/− mice. Nat Biotechnol. 2007 Aug;25(8):903–10.

23. Ponnudurai, T., Leeuwenberg, A., & Meuwissen, J. (n.d.). Chloroquine sensitivity of isolates of Plasmodium falciparum adapted to in vitro culture. Tropical and Geographical Medicine., 33(1), 50–54.

24. Vaughan AM, Mikolajczak SA, Camargo N, Lakshmanan V, Kennedy M, Lindner SE, Miller JL, Hume JCC, Kappe SHI. A transgenic Plasmodium falciparum NF54 strain that expresses GFP–luciferase throughout the parasite life cycle. Molecular and Biochemical Parasitology. 2012 Dec;186(2):143–7.

25. Research and Reference Reagent Resource Center (MR4). http://www.mr4.org.

26. Trager W, Jensen JB. Human malaria parasites in continuous culture. Science. 1976;193(4254):673–675. doi:10.1126/science.781840.

27. Feldmann AM, Ponnudurai T. Selection of Anopheles stephensi for refractoriness and susceptibility to Plasmodium falciparum. Med Vet Entomol. 1989;3(1):41–52. doi:10.1111/j.1365-2915.1989.tb00473.x.

28. Pewkliang Y, Rungin S, Lerdpanyangam K, Duangmanee A, Kanjanasirirat P, Suthivanich P, Sa-ngiamsuntorn K, Borwornpinyo S, Sattabongkot J, Patrapuvich R, Hongeng S. A novel immortalized hepatocyte-like cell line (imHC) supports in vitro liver stage development of the human malarial parasite Plasmodium vivax. Malar J. 2018 Dec;17(1):50.

29. Usui M, Fukumoto S, Inoue N, Kawazu S ichiro. Improvement of the observational method for Plasmodium berghei oocysts in the midgut of mosquitoes. Parasites Vectors. 2011;4(1):118.

30. Sack BK, Mikolajczak SA, Fishbaugher M, Vaughan AM, Flannery EL, Nguyen T, Betz W, Jane Navarro M, Foquet L, Steel RWJ, Billman ZP, Murphy SC, Hoffman SL, Chakravarty S, Sim BKL, Behet M, Reuling IJ, Walk J, Scholzen A, Sauerwein RW, Ishizuka AS, Flynn B, Seder RA, Kappe SHI. Humoral protection against mosquito bite-transmitted Plasmodium falciparum infection in humanized mice. npj Vaccines. 2017 Dec;2(1):27.

31. Kennedy M, Fishbaugher ME, Vaughan AM, Patrapuvich R, Boonhok R, Yimamnuaychok N, Rezakhani N, Metzger P, Ponpuak M, Sattabongkot J, Kappe SH, Hume JC, Lindner SE. A rapid and scalable density gradient purification method for Plasmodium sporozoites. Malar J. 2012;11(1):421.

32. Lupton EJ, Roth A, Patrapuvich R, Maher SP, Singh N, Sattabongkot J, Adams JH. Enhancing longevity of Plasmodium vivax and P. falciparum sporozoites after dissection from mosquito salivary glands. Parasitology International. 2015 Apr;64(2):211–8.

33. Angulo-Barturen I, Jiménez-Díaz MB, Mulet T, Rullas J, Herreros E, Ferrer S, Jiménez E, Mendoza A, Regadera J, Rosenthal PJ, Bathurst I, Pompliano DL, Gómez de las Heras F, Gargallo-Viola D. A Murine Model of falciparum-Malaria by In Vivo Selection of Competent Strains in Non-Myelodepleted Mice Engrafted with Human Erythrocytes. Keiser J, editor. PLoS ONE. 2008 May 21;3(5):e2252.

34. Golde WT, Gollobin P, Rodriguez LL. A rapid, simple, and humane method for submandibular bleeding of mice using a lancet. Lab Anim. 2005 Oct;34(9):39–43.

35. Lamikanra AA, Dobaño C, Jiménez A, Nhabomba A, Tsang HP, Guinovart C, Manaca MN, Quinto L, Aguilar R, Cisteró P, Alonso PL, Roberts DJ, Mayor A. A direct comparison of real time PCR on plasma and blood to detect Plasmodium falciparum infection in children. Malar J. 2012 Dec;11(1):201.

36. Shi B, Abrams M, Sepp-Lorenzino L. Expression of Asialoglycoprotein Receptor 1 in Human Hepatocellular Carcinoma. J Histochem Cytochem. 2013 Dec;61(12):901–9.

37. Westerink WMA, Schoonen WGEJ. Cytochrome P450 enzyme levels in HepG2 cells and cryopreserved primary human hepatocytes and their induction in HepG2 cells. Toxicology in Vitro. 2007 Dec;21(8):1581–91.

38. Hermsen CC, Telgt DSC, Linders EHP, van de Locht LATF, Eling WMC, Mensink EJBM, Sauerwein RW. Detection of Plasmodium falciparum malaria parasites in vivo by real-time quantitative PCR. Molecular and Biochemical Parasitology. 2001 Dec;118(2):247–51.

39. Shokoples SE, Ndao M, Kowalewska-Grochowska K, Yanow SK. Multiplexed Real-Time PCR Assay for Discrimination of Plasmodium Species with Improved Sensitivity for Mixed Infections. Journal of Clinical Microbiology. 2009 Apr 1;47(4):975–80.

40. Wampfler R, Mwingira F, Javati S, Robinson L, Betuela I, Siba P, Beck HP, Mueller I, Felger I. Strategies for Detection of Plasmodium species Gametocytes. Paul RE, editor. PLoS ONE. 2013 Sep 27;8(9):e76316.

41. Franke-Fayard B, Trueman H, Ramesar J, Mendoza J, van der Keur M, van der Linden R, Sinden RE, Waters AP, Janse CJ. A Plasmodium berghei reference line that constitutively expresses GFP at a high level throughout the complete life cycle. Molecular and Biochemical Parasitology. 2004 Sep;137(1):23–33.

42. Bon C, Hofer T, Bousquet-Mélou A, Davies MR, Krippendorff BF. Capacity limits of asialoglycoprotein receptor-mediated liver targeting. mAbs. 2017 Nov 17;9(8):1360–9.

43. Pybus BS, Marcsisin SR, Jin X, Deye G, Sousa JC, Li Q, Caridha D, Zeng Q, Reichard GA, Ockenhouse C, Bennett J, Walker LA, Ohrt C, Melendez V. The metabolism of primaquine to its active metabolite is dependent on CYP 2D6. Malar J. 2013 Dec;12(1):212.

44. Kaushansky A, Douglass AN, Arang N, Vigdorovich V, Dambrauskas N, Kain HS, Austin LS, Sather DN, Kappe SHI. Malaria parasites target the hepatocyte receptor EphA2 for successful host infection. Science. 2015 Nov 27;350(6264):1089–92.

45. Silvie O. Cholesterol contributes to the organization of tetraspanin-enriched microdomains and to CD81-dependent infection by malaria sporozoites. Journal of Cell Science. 2006 May 15;119(10):1992–2002.

46. Silvie O, Rubinstein E, Franetich JF, Prenant M, Belnoue E, Rénia L, Hannoun L, Eling W, Levy S, Boucheix C, Mazier D. Hepatocyte CD81 is required for Plasmodium falciparum and Plasmodium yoelii sporozoite infectivity. Nat Med. 2003 Jan;9(1):93–6.

47. Foquet L, Hermsen CC, Verhoye L, van Gemert GJ, Cortese R, Nicosia A, Sauerwein RW, Leroux-Roels G, Meuleman P. Anti-CD81 but not anti-SR-BI blocks Plasmodium falciparum liver infection in a humanized mouse model. Journal of Antimicrobial Chemotherapy. 2015 Feb 4;dkv019.

48. Risco-Castillo V, Topçu S, Son O, Briquet S, Manzoni G, Silvie O. CD81 is required for rhoptry discharge during host cell invasion by *P lasmodium yoelii* sporozoites: CD81 and *Plasmodium* sporozoite rhoptry discharge. Cell Microbiol. 2014 Oct;16(10):1533–48.

49. Ng S, March S, Galstian A, Gural N, Stevens KR, Mota MM, Bhatia SN. Towards a Humanized Mouse Model of Liver Stage Malaria Using Ectopic Artificial Livers. Sci Rep. 2017 Jun;7(1):45424.

50. Morosan S, Battaglia S, Blanc C, Eling W, Sauerwein R, Hannoun L, Belghiti J, Brechot C, Kremsdorf D, Druilhe P. Liver-Stage Development of Plasmodium falciparum, in a Humanized Mouse Model. Journal of Infectious Diseases. 2005 Nov;193:996–1004.

51. Marcsisin SR, Reichard G, Pybus BS. Primaquine pharmacology in the context of CYP 2D6 pharmacogenomics: Current state of the art. Pharmacology & Therapeutics. 2016 May;161:1–10.

52. St Jean PL, Xue Z, Carter N, Koh GCKW, Duparc S, Taylor M, Beaumont C, Llanos-Cuentas A, Rueangweerayut R, Krudsood S, Green JA, Rubio JP. Tafenoquine treatment of Plasmodium vivax malaria: suggestive evidence that CYP2D6 reduced metabolism is not associated with relapse in the Phase 2b DETECTIVE trial. Malar J. 2016 Dec;15(1):97.

53. D’Souza AA, Devarajan PV. Asialoglycoprotein receptor mediated hepatocyte targeting — Strategies and applications. Journal of Controlled Release. 2015 Apr;203:126–39.

54. Flannery EL, Foquet L, Chuenchob V, Fishbaugher M, Billman Z, Navarro MJ, Betz W, Olsen TM, Lee J, Camargo N, Nguyen T, Schafer C, Sack BK, Wilson EM, Saunders J, Bial J, Campo B, Charman SA, Murphy SC, Phillips MA, Kappe SHI, Mikolajczak SA. Assessing drug efficacy against Plasmodium falciparum liver stages in vivo. JCI Insight. 2018 Jan 11;3(1):e92587.

55. Douglass AN, Kain HS, Abdullahi M, Arang N, Austin LS, Mikolajczak SA, Billman ZP, Hume JCC, Murphy SC, Kappe SHI, Kaushansky A. Host-based Prophylaxis Successfully Targets Liver Stage Malaria Parasites. Molecular Therapy. 2015 May;23(5):857–65.

56. Moreno A, Ferrer E, Arahuetes S, Eguiluz C, Rooijen NV, Benito A. The course of infections and pathology in immunomodulated NOD/LtSz-SCID mice inoculated with Plasmodium falciparum laboratory lines and clinical isolates. International Journal for Parasitology. 2006 Mar;36(3):361–9.

57. March S, Ng S, Velmurugan S, Galstian A, Shan J, Logan DJ, Carpenter AE, Thomas D, Sim BKL, Mota MM, Hoffman SL, Bhatia SN. A Microscale Human Liver Platform that Supports the Hepatic Stages of Plasmodium falciparum and vivax. Cell Host & Microbe. 2013 Jul;14(1):104–15.

